# Presence of sodium taurocholate co-transporting polypeptide and Hepatitis B replication marker on placenta: Another home for the virus

**DOI:** 10.1101/2022.01.05.475013

**Authors:** Garima Garg, Meenu MN, Kajal Patel, Shashank Purwar, Sramana Mukhopadhyay, Nitu Mishra, Sudheer Gupta, Sumit Kumar Rawat, Ritu Khosla, Jitendra Singh, Shashwati Nema, Debasis Biswas, Anirudh K Singh, Ashish Kumar Vyas

## Abstract

**Background:** The role of sodium taurocholate co-transporting polypeptide (NTCP), in facilitating the binding of Hepatitis B virus (HBV) on surface of hepatocytes is well documented. Expression of NTCP in extra hepatic cells may make these cells susceptible to HBV infection and support cellular proliferation akin to hepatocytes. *Placental replication of HBV is not well explored*. In this study we have assessed the expression of NTCP and HBV replication markers (HBeAg, HBcAg, and HBV DNA) in placental cells, to investigate if these cells act as host for HBV.

**Methods:** Fourty one HBsAg+ve pregnant women along with 10 healthy controls were enrolled after obtaining informed consent. The HBV DNA in placenta was detected by qPCR using primers for X and core ORF. Expression of NTCP in placenta was analyzed by qRT-PCR and further investigated by immunohistochemistry (IHC) along with HBV replication biomarkers, HBeAg, and HBcAg.

**Results:** HBsAg positive subjects were divided in two groups on the basis of viral load [High Viral Load (HVL) Group; viral load ≥ 2000IU/ml, Low Viral Load (LVL) Group; viral load <2000IU/ml] according to INASL guidelines 2018. HBV infected females showed increased expression of NTCP in trophoblasts of placenta compared to control group (HVL 3.69±0.13 Vs Control 1.74±0.15, p=0.0117). Furthermore, significant difference in NTCP expression was also observed between HVL and LVL group (HVL 3.69±0.13 Vs LVL 1.98±0.17, p=0.022) and positively correlated with the maternal HBV DNA load. Membranous and/or cytoplasmic immunostaining of NTCP, and cytoplasmic staining of HBeAg and HBcAg in trophoblasts along with presence of HBV DNA indicated that trophoblasts are not only susceptible to HBV infection but may also be a site for viral replication.

**Conclusions:** This is the pioneer study, which demonstrates expression of NTCP on placenta which may facilitate the entry of HBV. Furthermore, the study establishes the presence of HBeAg in placenta of patients without circulating HBeAg, indicating these cells may act as replication host/reservoir. This pioneering finding hints at the possibility of exploring the potential of NTCP blocking strategies in preventing vertical transmission of HBV.

## Introduction

Chronic Hepatitis B (CHB) infection is a major global health problem with 257 million infected people worldwide resulting in 887000 deaths annually [1 ]. Vertical transmission of HBV is a leading cause of chronicity^2^. About 90-95% children acquiring infection from mother develop chronicity compared to adults where this risk is just 5-10%. vertical transmission is main route of infection in developing countries. In India, 1-4% of population is chronic carrier of HBV. Patients with CHB have high risk of developing end-stage cirrhosis and hepatocellular carcinoma[3]. Maternal hepatitis B e antigen (HBeAg) and high viral load are associated with HBV transmission to their newborns [2,4]. Earlier studies suggest that among the three mode of vertical transmission (Intrauterine or prenatal, Natal and Postnatal) [5]. However, precise mechanism of trans-placental routes of infection and vertical transmission of mother to baby are understudied. In one of the few available studies, we recently determined the role of maternal immunity in mother to baby vertical transmission[6] and the increased expression of HBV entry co-receptor Asialoglycoprotein receptor (ASGPR) in placental cells of transmitting mothers [7].

Sodium Taurocholate Co-transporting Polypeptide (NTCP) is a functional receptor of HBV. NTCP is an integral membrane glycoprotein involved in uptake of glycine/taurine-conjugated bile acids. It is considered to be exclusively expressed on hepatocytes [8]. Presence of NTCP is a key determinant of hepatotropism of HBV but its presence on placental cells is largely unknown and if present, it may play a crucial role in vertical transmission. Studies in primary trophoblast culture showed susceptibility of trophoblasts for HBV infection but there has been no such study on clinical specimens of placenta.

Previously serological studies showed that HBeAg can cross the placenta [9] but whether replication occurs in placenta was not explored. After getting entry through NTCP, does HBV also replicate inside trophoblasts is a considerable question.

In the present study, we not only found the expression of NTCP on placental trophoblasts, but also observed the presence of HBeAg and HBV DNA (both markers of active HBV replication) in trophoblast cells. Thus, through demonstration of both HBV receptor and replication in placental tissues, this study generates novel evidence for the placenta acting as a target and reservoir organ for this virus and provides a mechanistic basis for vertical transmission.

## Results

### 2.1 Virological and serological characteristics

The demographic and biochemical profile of subjects enrolled in the study is shown in **Table 1**. We did not find difference in base line clinical parameter between the HVL and LVL groups. Serum HBeAg analysis in HVL group showed 26.66 % and none were positive in LVL group.

### 2.2 Presence of HBV infection in placenta

Presence of HBsAg and HBcAg in placenta denotes the infection of HBV in placental cells [9,11]. In present study we confirmed the HBV infection in placental cells by determining HBcAg by IHC. HBcAg was present in more than 70% samples of HVL group and approximately 80% samples of LVL group (Figure 1 A). Control placentas were also examined for any nonspecific binding. We did not find any significant difference in the IHC score of HBcAg in HVL and LVL groups (HVL 94.62±9.46 Vs LVL 76.5±8.68 p = 0.9253) Figure 1 B).

| Parameter | Control (n=10) | LVL(n=19) | HVL(n=22) |
| --- | --- | --- | --- |
| Age years | 23.9±3.755 | 22.47±2.54 | 24.51±3.17 |
| HBV DNA<br>(IU/ml) | -- | 39.2<br>(3.58-1790) | 3.0*10<sup>6</sup><br>(2.2*12<sup>3</sup>-4.78 x 10<sup>8</sup>) |
| AST (IU) | 19.9 ±15.75 | 99.81±67.57 | 105.6±95.85 |
| ALT (IU) | 29.51±12.24 | 78.38 ±37.48 | 56.71 ±34.28 |
| HBeAg | -- | NIL | 26.66% |

**Figure 1.**
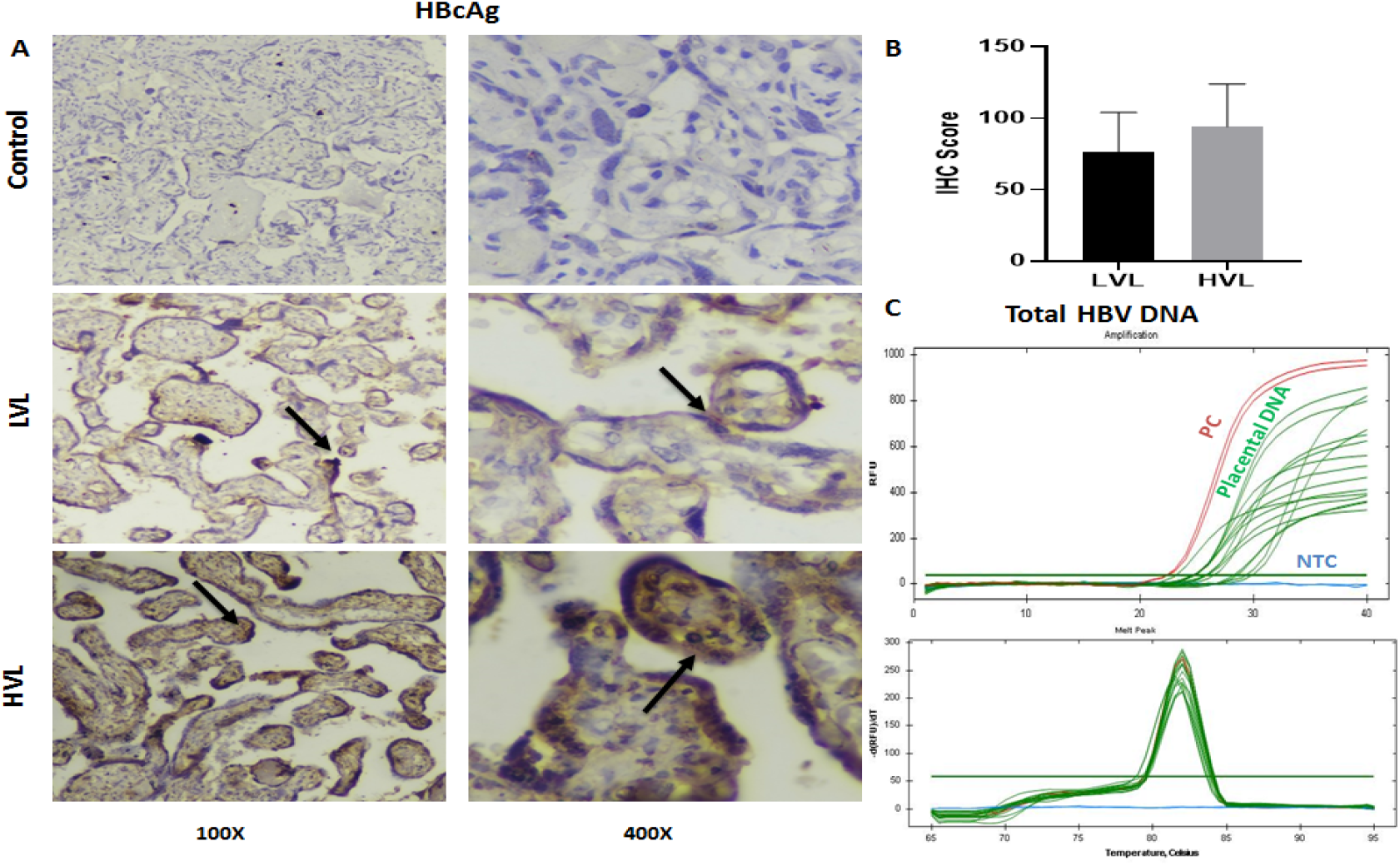
Confirmation of Hepatitis B Virus infection in placenta. Representative Figure showed the presence of HBcAg in placenta through Immunohistochemistry (IHC), cytoplasmic staining in LVL and HVL groups was observed, and no staining in control **(A)**. IHC Scores of HVL and LVL groups showed no significant difference (HVL 71±7.22 Vs LVL 65.5±7.43 p = 0.8687). Significance was considered p-value was calculated for a risk threshold *α*=0.05, p* <0.0332 (**B**). Detection of total HBV DNA in placenta through qPCR Amplification curve and single peak in melt curve **(C)**.

HVL- High Viral Load Group, LVL- Low Viral Load Group, PC- Positive Control, NTC- No Template Control As presence of HBV DNA is also considered as a marker of infection, we performed qPCR for X and Core ORF to detect HBV DNA was detected in placental samples (supplementary Figure 1) from both groups and found that 92.30% of HVL and 62.5% of LVL samples gave positive amplification for these targets (Figure 1 C). The high percentage of HBV DNA positivity in both groups and stringent positivity analysis through HBcAg indicated placental infection in most of the samples. Our next aim wais to investigate whether HBV entry receptor NTCP expresses on placental cells.

### 2.3 NTCP expresses on placental cells

Encouraged by finding markers for HBV infection in placenta, we postulated that placenta may act as reservoir and facilitate viral replication. NTCP is the primary receptor for virus on hepatocytes and is thought to be exclusive to the cells. However, for placental cells to act as reservoir, these cells must express NTCP receptor. We started with probing the expression of receptor in placenta. The expression of NTCP was searched in the data submitted for different cell types/tissues to the “The Human Protein Atlas” (www.proteinatlas.org). The consensus normalized expression for single cell RNA (nTPM) from all single cell types have been shown in Figure 2.

**Figure 2.**
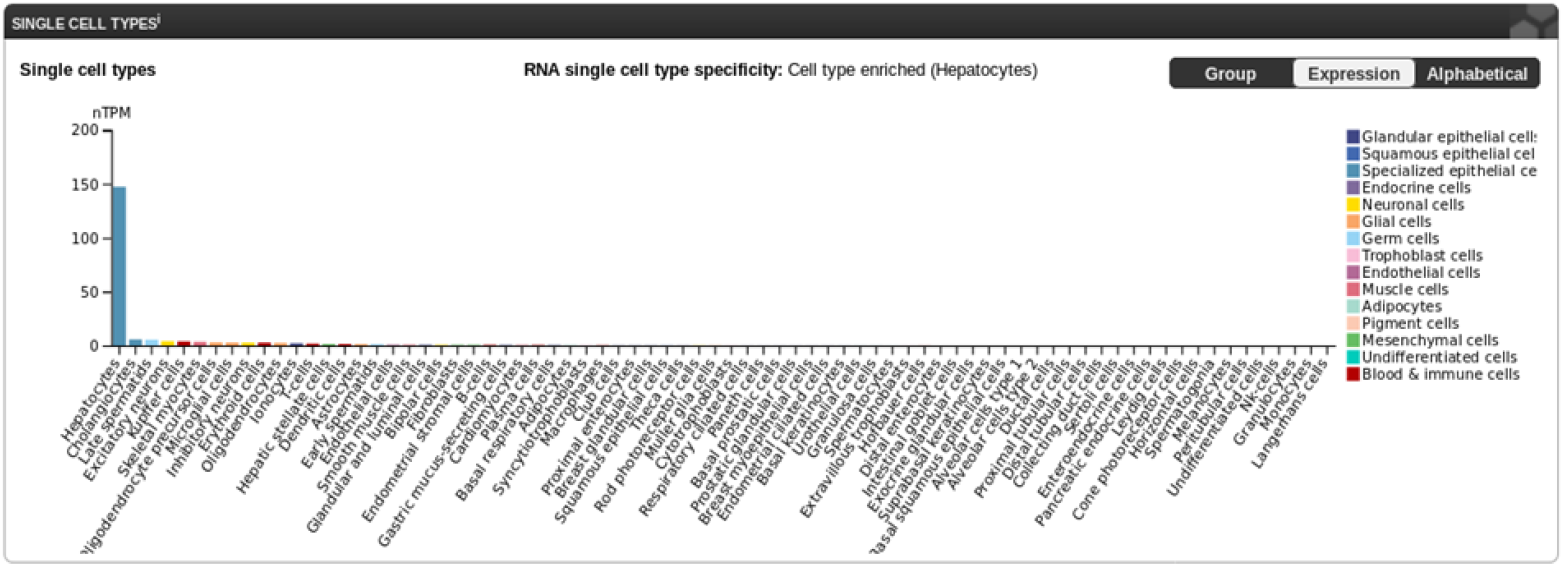
Plot showing the consensus normalized expression of NTCP in different cell types. Color-coding in the plot, is based on cell type groups, each consisting of cell types with functional features in common.

It can be observed that the expression of NTCP in healthy human cell types is reported mainly in Hepatocytes and to a lower level in Cholangiocytes, Late spermatids, Excitatory neurons, Kupffer cells, Skeletal myocytes, Oligodendrocyte precursor cells, Microglial cells, Erythroid cells, Inhibitory neurons, Oligodendrocytes, Ionocytes, T-cells, Dendritic cells, Hepatic stellate cells, Astrocytes, Early spermatids, Endothelial cells, Smooth muscle cells & B-cells. The cells specific to placenta have not been reported to express NTCP in the studies available in “The Human Protein Atlas”.

We started examining expression of NTCP in our placental samples and interestingly found that NTCP is expressed in placenta as well (Figure 3). <u>Quantification of NTCP expression was done by qRT-PCR.</u> While NTCP expression in LVL group was comparable to the control group (LVL 1.98±0.17 Vs Control 1.74±0.15, p= 0.9213; *<0.0332), it was significantly upregulated in HVL group when compared with control group (HVL 3.69±0.13 Vs Control 1.74±0.15, p=0.0117; *<0.0332) or LVL group (HVL 3.69±0.13 Vs LVL 1.98±0.17, p=0.022; *<0.0332). This suggest that NTCP expression might be associated with viral load (Figure 3 A). To further ensure NTCP is translated we performed <u>Immunohistochemistry</u> IHC using NTCP specific antibody. Membranous and / or cytoplasmic staining was considered positive with reference to liver as positive control. Expression of NTCP was observed mainly in trophoblasts while stromal and endothelial cells were not stained (Figure 3 B). Membranous expression was largely observed in HVL group placenta. For confirmation of specificity of NTCP antibodies we also analyzed the IHC of other tissues (Breast tissue, Myometrium and Endometrium of uterus) for NTCP expression. (Supplementary Figure 2). As expected, we did not find expression of NTCP on breast tissue and layers of uterus, circumvents postnatal and prenatal risk, respectively. There was a significant difference between HVL and control group (HVL 185.5±9.71 Vs Control 65.5±5.4, p = 0.011; *<0.0332). Whereas, no significant difference was obtained between HVL and LVL groups (HVL 185.5±9.71 Vs LVL 130.5±9.62, p = 0.333) and LVL and control group (LVL 130.5±9.62 Vs Control 65.5±5.4, p = 0.221) (Figure 3 C).

**Figure 3.**
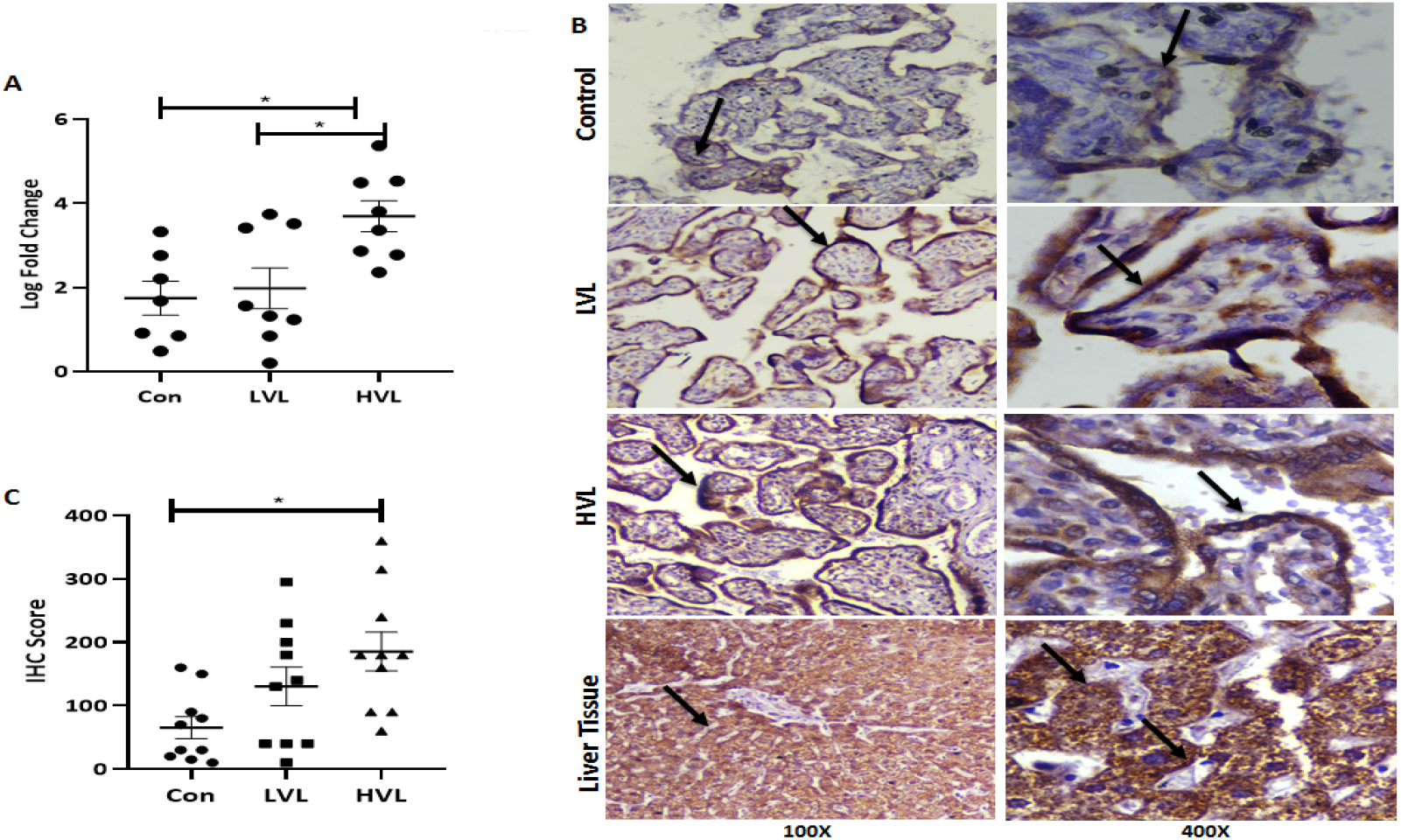
Expression of NTCP in placenta. The log fold change in expression of NTCP in placenta among the groups **(A)**. Representative Figure of expression of NTCP in placenta through Immunohistochemistry, weak cytoplasmic staining in control, moderate to strong membranous and cytoplasmic staining in LVL and HVL group was observed. Strong membranous and cytoplasmic staining of non pathological liver tissue was served as positive control **(B)**. IHC Scores were plotted against log of HBV DNA viral load (IU/ml) **(C)**. Con- Control, HVL- High Viral Load Group, LVL- Low Viral Load Group

Presence of NTCP on placental cells indicate another home for HBV to validate this point we have assess the markers of HBV replication on placental cells.

### 2.4 Presence of marker of HBV replication in placental cells

In order to determine if HBV can replicate in placental cells, we probed for the presence of HBeAg, a marker of viral replication, in trophoblasts using IHC. Interestingly, we found the presence of HBeAg in trophoblast cells. We observed cytoplasmic staining in the 40% HVL group and 80% LVL group while there was no staining in the control group placentas showing specific immunostaining (Figure. 4 A).

**Figure 4.**
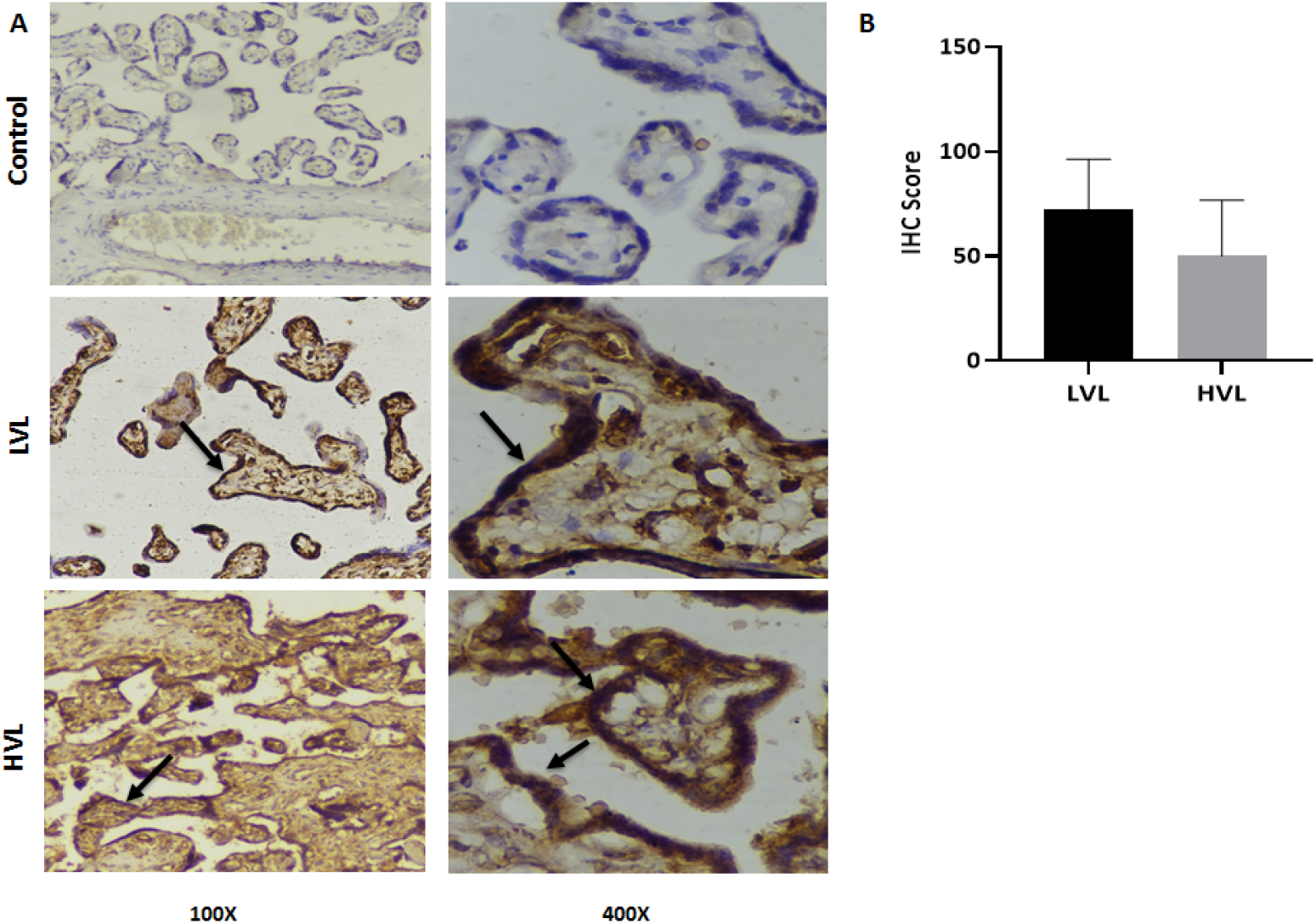
Presence of HBeAg in placenta. Representative Figure of detection of HBeAg in placenta through Immunohistochemistry, no staining in control and strong cytoplasmic staining in LVL and HVL groups was observed **(A)**. IHC Scores of HVL and LVL between the groups (**B)**. Two-tailed p-value was calculated for a risk threshold *α*=0.05. p* <0.0332.

<u>To eliminate any false positivity, we did not consider weak intensity staining as positive.</u> On comparison of percentage of samples positive for peripheral and/or placental HBeAg, we found only 7.14% was positive for both while 14.28% were positive for HBeAg in circulation but not for placenta showing not all the peripheral positivity turned to placental. <u>Surprisingly in 50% cases we found placental positivity but not peripheral affirming our finding that HBeAg is placenta derived as a result of active replication and not infiltered from periphery</u> and 28.57% was negative for both. We did not find significant difference between the presence of HBeAg in HVL and LVL groups (HVL 50±8.43 Vs LVL 72.5±7.51 p = 0.2204; *<0.0332) (Figure 4 B). Our finding strongly indicates that HBV can also replicate in placenta.

### 2.5 Association of peripheral viral load with placental HBV infection markers and NTCP

In order to analyse the correlation between placental infection markers (i.e., HBcAg and HBeAg) and NTCP with viral load, we found a significant positive correlation between log fold change of NTCP expression with viral load (PC = 0.7118, R^2^ = 0.3796, p = 0.0027**; p**0.0021) although correlation was not significant when we compared IHC scores of NTCP (PC = 0.3577, R^2^ = 0.0518) indicating post transcriptional modulation which has to be assessed on larger sample size. We also did correlation analysis with peripheral viral load with IHC score of HBcAg (PC = 0.6160, R^2^ = 0.0408), and HBeAg (PC = -0.05213, R^2^ = 0.0009) (Figure 5 A-C) but no correlation was found.

**Figure 5.**
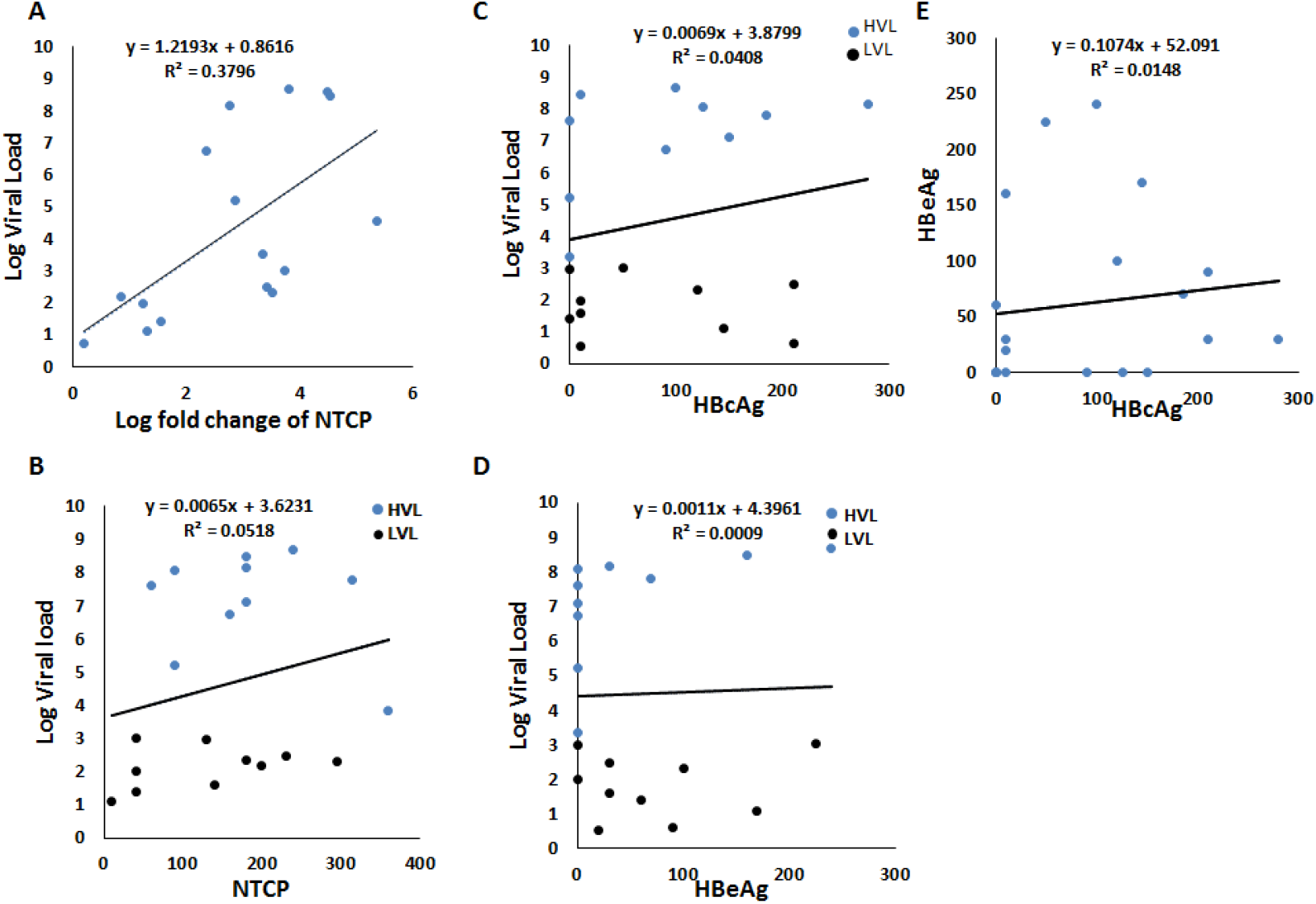
Correlation of Peripheral viral load with IHC score of NTCP, HBcAg and HBeAg. Dot plots showing correlation between log fold change of NTCP expression load (PC = 0.7118, R^2^ = 0.3796, p = 0.0027**; p**0.0021) **(A)** IHC score of NTCP (PC = 0.3577, R^2^ = 0.0518) **(B), IHC score of** HbcAg (PC = 0.04907, R^2^ = 0.0139) **(C)** IHC score of HBeAg (PC = -0.05213, R^2^ = 0.0009) **(D)**, with peripheral viral load. Correlation coefficient was calculated using Spearman’s correlation test. **(E)** Correlation of IHC score of HBeAg with IHC score of HBcAg in placental tissues (PC = 0.4015, R^2^ =0.0112).

We further analyze the association between IHC score of HBeAg and HBcAg in the placental samples but did not find statistically significant association between them (PC = 0.1153, R^2^ =0.0148) (Figure 5 D and Supplementary Figure 3).

## Discussion

As HBV vertical transmission is the primary cause of chronic HBV infection worldwide[12], inhibiting it is the most effective approach towards eradicating chronic HBV infection. Infants infected with HBV face the risk of severe complications due to genetic, epigenetic, and environmental factors. In spite of universal vaccination program, 5% or more of vaccinated newborns still do not achieve protective levels of anti-hepatitis B virus surface antigen titers [13]. The precise mechanism behind HBV vertical transmission is not known. Our previous study showed presence of ASGPR on placenta and circulating dendritic cells which facilitates the endocytosis of HBV^7^. Paganelli et al., also found presence of ASGPR on umbilical cord matrix stem cells (UCMS) [14], and showed its competitive inhibition prevents binding and uptake of HBV. In order to establish infection, HBV must bind to its specific receptor. NTCP is a primary receptor of HBV. It was known to present only on hepatocytes and is responsible for the hepatotropism of HBV. It’s presence on non-hepatic cells is not well known. Presence of NTCP is critical to make in vitro culture systems susceptible for HBV infection [15]. Indicating that its presence on non-hepatic tissues can make them susceptible towards infection. Here, we far the first time report the presence of NTCP on placenta along with HBV replication markers. HBeAg is a marker of active replication. Previous studies suggested that HBeAg can cross the placenta but it was only based on the serological findings of mother and newborn. In our present study we not only observed the presence of HBeAg in the placental cells / trophoblasts but also in 50% cases who were negative for HBeAg in circulation were positive in placenta along with HBV DNA strongly suggests that placenta can also serve as a host tissue for HBV replication.

We understand that this novel finding provides a mechanistic basis for vertical transmission of HBV and also opens up potential strategies for arresting vertical transmission targeted at NTCP blocking by treatment with Myrcludex B and/or through other alternative strategies may be used for therapeutic intervention of vertical transmission.

Our study suffers from the limitation of small sample size and the inability to experimentally demonstrate the prevention of HBV infection of placental tissues through blockage of NTCP receptor.

## Materials and Method

### HBV pregnant females

A total of 27407 pregnant females were screened for HBsAg during their antenatal routine check-up in the in a tertiary care teaching hospital in central India over a period of 2 years from 2020-2021. A total of 432 of pregnant females were found positive for HBsAg (1.57%) and negative for hepatitis A, hepatitis C, hepatitis E and human immunodeficiency virus (HIV). None of the pregnant females had any systemic illness, autoimmune disease or inherited metabolic disorder. Out of 432, we were able to follow 59 HBsAg+ve pregnant females, till delivery due to loss of follow ups, lack of consent and other complications. Out of 59, 22 had HBV DNA ≥2000IU/ml, 19 had HBV DNA <2000IU/ml and in 18 samples HBV DNA was below the detection limit. A total 41 HBsAg+ve and 10 healthy pregnant females were included in this study.

### Study Groups

#### HBsAg Positive mothers (n=41)

HBsAg positive subjects were divided in two groups on the basis of viral load according to INASL guidelines 2018 [10]. Pregnant females, had HBV DNA load ≥ 2000IU/ml were categorized as <u>High viral load (HVL)</u> group (n= 22) and pregnant females, who had HBV DNA load <2000IU/ml were categorized as <u>Low viral load (LVL)</u> group (n=19).

#### Control Group (n=10)

Pregnant females, who were negative for HBsAg and gave birth to healthy newborns, were identified as healthy and were included in the study as controls.

The Informed consent was obtained from each participant before enrolment in the study. The clinical and biochemical assessment of the subjects was done according to the study protocol. None of the participants was receiving anti-HBV drugs.

### Sampling of serological and virologic studies

At term, 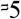 ml of peripheral blood sample was collected in EDTA tubes and placental tissues were obtained from maternal decidua (within 6 hours) of delivery. The plasma was separated within an hour of blood collection and immediately stored at -80°C till further use.

#### Peripheral HBV DNA Quantification

HBV DNA quantitation was done with 500 μl plasma using COBAS Taq Man HBV test with high purity extraction (Roche Diagnostics, USA), as per the manufacturer’s protocol. This real-time PCR assay is based on dual labelled hybridization probe targeting the pre-core and core regions. Results were expressed as IU/ml. The lower limit of detection for the assay was <6 IU/ml.

### DNA and RNA Extraction from placental tissues

DNA and RNA was extracted from placental tissues frozen in RNA Liv (Himedia). DNA from approximately 20 mg placental tissue was extracted using QIAmp Blood DNA mini kit, Qiagen, Germany using manufacturer’s protocol. Approximately 20 mg placental tissue was homogenised using TissueLyser LT (Qiagen) and total RNA was extracted using QIAamp®RNA Blood Mini kit, Qiagen, Germany according to manufacturer’s protocol. DNA and RNA was quantified using NanoDrop^™^ Spectrophotometer (Thermo Scientific^™^). High-Capacity cDNA Reverse Transcription Kit (Applied Biosystems, Vilnius, LT) was used for complementary DNA synthesis using 1µg of RNA.

### Detection of HBV DNA in placenta

qPCR was performed with PowerUp™ SYBR™ Green Master Mix kit, (Applied Biosystems, Vilnius, LT) using an BioRad CFX96 Real Time PCR System (BioRad) with primers specific for total HBV DNA targeting X and core ORF (HBV F: GAGGCTGTAGGCATAAATTGGTC, HBV R: AACTCCACAGAAGCTCCAAATTC). Every sample was independently analysed in triplicates.

### Detection of HBcAg and HBeAg in placenta by Immunohistochemistry (IHC)

Five-micron thick sections were cut from Formalin Fixed Paraffin Embedded (FFPE) tissue samples and deparaffinized by xylene followed by heat induced antigen retrieval by boiling at 100 °C for 10 minutes in citrate buffer (pH= 6). Tissue sections were incubated with the following primary antibodies overnight at 4°C: HBcAg (bsm-2000M; Bioss, 1:25), HBeAg (bsm-2022M; Bioss, 1:25). Polyexcel HRP/DAB detection system, PathnSitu (USA) was used for staining using manufacturer’s protocol and counterstaining was done with haematoxylin. Images were captured using Nikon DS-Ri2 camera on a Nikon Eclipse Ci microscope.

### Quantification of NTCP in placental tissues using qRT-PCR

cDNA prepared from placental RNA was subjected to qPCR using PowerUp™ SYBR™ Green Master Mix kit (Applied Biosystems, Vilnius, LT) on BioRad CFX96 Real Time PCR System (BioRad). The specific exon spanning primers of NTCP were designed (NTCP Fwd: CTTCTGCCTCAATGGACGGT; NTCP Rv: GCCACATTGAGGATGGTGGA). qRT-PCR of every sample was done in triplicates to normalize NTCP expression level; GAPDH was used as reference gene. Subsequently, the relative gene expression values were determined using log of 2^-ΔΔCT^ and specificity of product was assured by single melt peak.

### Detection of NTCP in placenta by Immunohistochemistry (IHC)

Five-micron tissue sections were cut, and deparaffinized by xylene followed by heat induced antigen retrieval using boiling at 100° c for 10 minutes in Tris buffer (pH-9). Tissues were then incubated with the primary antibody anti-NTCP (BS1985R; Bioss, 1:1000) for 1 hour at room temperature, further staining was done using Polyexcel HRP/DAB detection system PathnSitu (USA) according to manufacturer’s protocol and were counterstained with haematoxylin. Healthy Liver section was taken as positive control. Breast, endometrial and myometrial tissues were also stained and analysed. Images were captured using Nikon DS-Ri2 camera on a Nikon Eclipse Ci microscope.

### IHC Quantification

IHC slides were score by a pathologist using single blinded strategy. Membranous and / or cytoplasmic staining was considered as positive. IHC score was calculated using formula; IHC score = percentage of positive cells * Score of Intensity of the staining. The Score of Intensity followed 1- Weak, 2- Mild, 3- Moderate and 4- Strong for NTCP and HBcAg and 0- Weak, 1- Mild, 2- Moderate and 3- Strong for HBeAg. IHC score was used to calculate statistical significance.

#### Statistical Analysis

All statistical tests were performed using GraphPad Prism for Windows version 8.0.1. Participant’s demographics are presented as median with range or mean with standard deviation. Categorical variables are presented as proportions while continuous variables are either presented as mean with standard error (SE) or median with range. Continuous variables were tested for normal distribution by Bartlett’s test followed by one way ANOVA (nonparametric) test and Tukey’s multiple comparison test to find the significance. Correlation was calculated by Spearman’s statistics. All statistical comparisons were two tailed and a p value of <0.0332 was considered statistically significant.

**Conclusion**, this study, first time demonstrating the presence of NTCP on placenta and in search of probable replication, we interestingly found the presence of HBV DNA and HBeAg in placental cells of HBeAg negative subjects in circulation. NTCP blocking by treatment with Myrcludex B and/or through other strategy may be used for therapeutic intervention of vertical transmission. Our findings are providing an insight on the probable new mechanism contributing in vertical transmission of the virus and expected to be useful for therapeutic approaches towards its prevention. In summary, this study provides an insight on the probable mechanism contributing to vertical transmission of the virus, which can be clinically exploited in future for its prevention and treatment.

## Author Contributions

Conceptualization, project administration and funding acquisition, Ashish Kumar Vyas; visualization and writing—original draft preparation, Garima Garg, methodology, data curation, and validation, Garima Garg, Meenu MN and Kajal Patel; formal analysis, Sramana Mukhopadhyay; resources, Nitu Mishra, Sumit Kumar Rawat; writing—review and editing, Anirudh K Singh, Ritu Khosla, Jitendra Singh and Shashwati Nema ; supervision and investigation, Shashank Purwar and Debasis Biswas;

## Funding

This research was funded by Department of Science and Technology, INDIA Grant No. DST/INSPIRE/04/2018/000019 and Department of Science and Technology, Science and Engineering Research Board Grant No CRG/2020/003856 to Dr. Ashish Kumar Vyas.

## Institutional Review Board Statement

The study was conducted in accordance with the Declaration of Helsinki, and approved by the Institutional Human Ethics Committee of All India Medical Sciences (AIIMS), Bhopal, India (Approval No. EF0110; date of approval 26.12.2019 and EF0232; date of approval 19.04.2021

## Informed Consent Statement

Informed consent was obtained from all subjects involved in the study

## Acknowledgments

We thank technical staff of Dept. of Pathology, AIIMS, Bhopal for help in IHC work. Nursing Staff of Sultania maternity hospital, Bhopal for participant enrolment. We thank to Ms. Priyal Gupta and Ms. Bijina John Mathew for their intellectual input.

## Conflicts of Interest

The authors declare no conflict of interest.

---

**Supplementary Figure 1.**
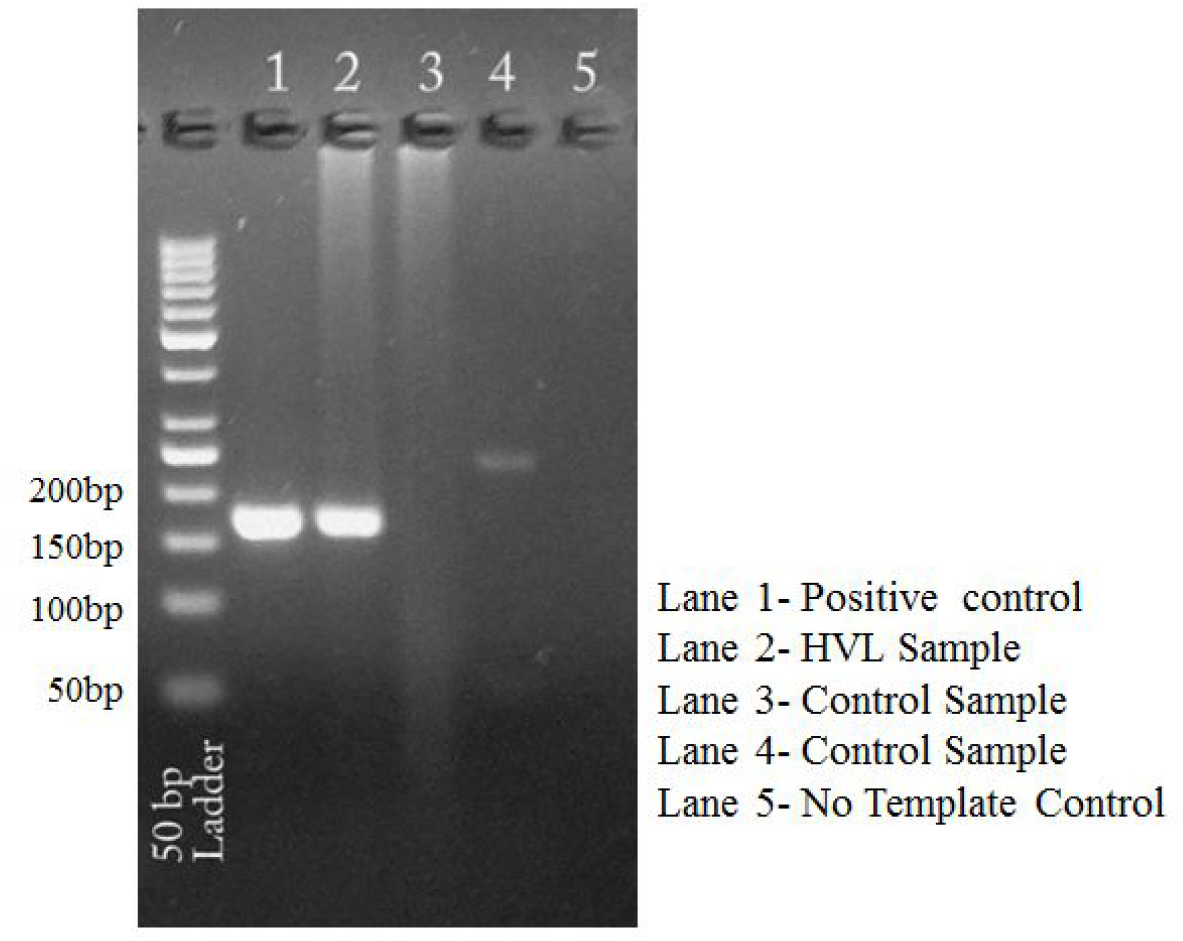
Gel image showing specific band of HBV DNA qPCR.

**Supplementary Figure 2.**
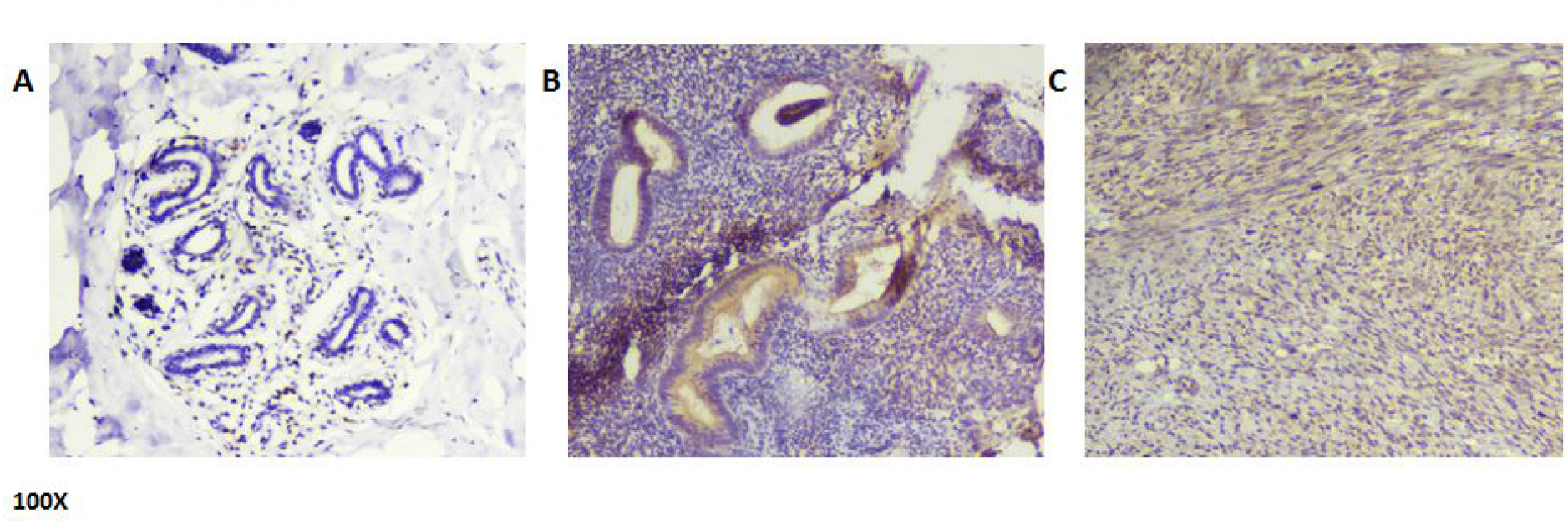
Expression of NTCP in different nonpathological tissues detected by Immunohistochemistry (IHC) (A) No staining was observed in Breast tissue, weak/ no staining in (B) Endometrium and (C) Myometrium

**Supplementary Figure 3.**
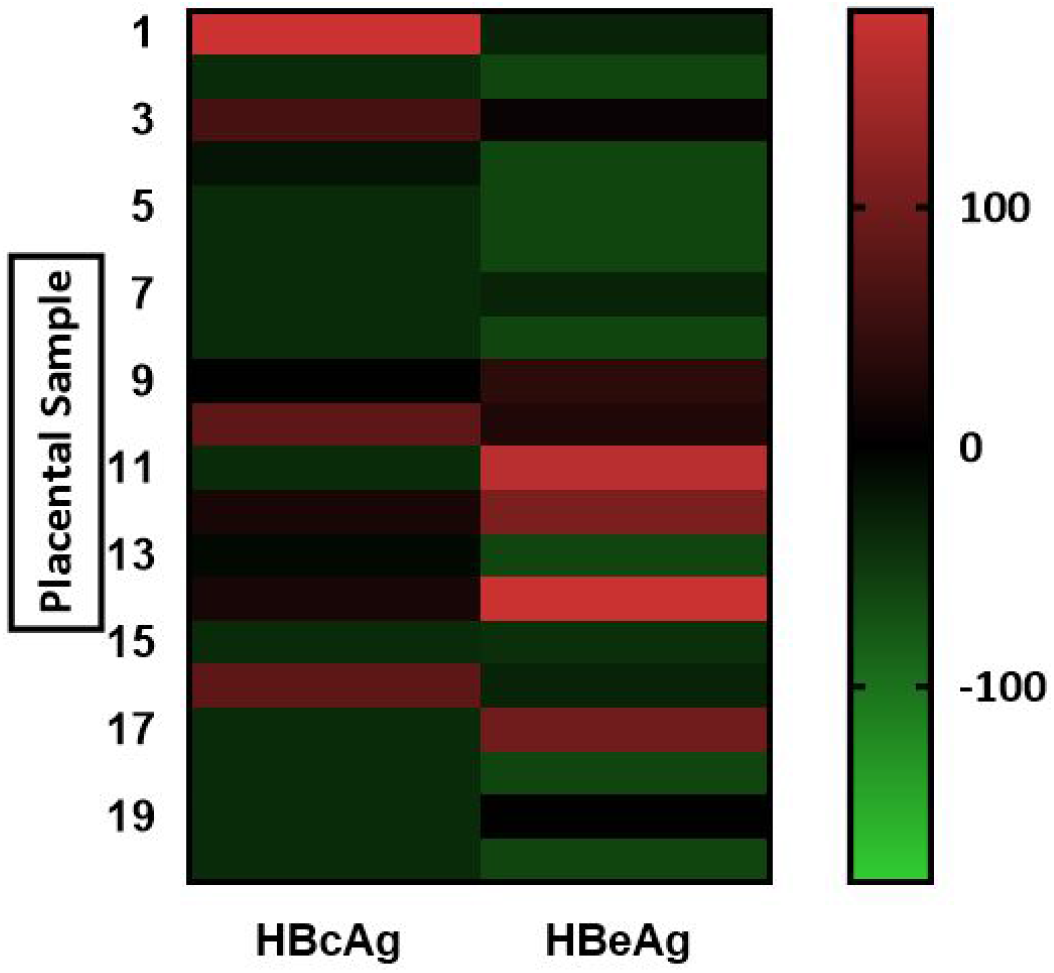
Heat Map showing sample wise comparison of HBcAg and HBeAg immunostaining

## Notes

### Competing Interest Statement

The authors have declared no competing interest.

